# Detecting bat environmental DNA from water-filled road-ruts in upland forest

**DOI:** 10.1101/2022.06.26.497664

**Authors:** Nathaniel T. Marshall, Daniel E. Symonds, Faith M Walker, Daniel E Sanchez, Zachary L. Couch, James D. Kiser

## Abstract

Declines in population and diversity of North American bats are rapidly occurring due to habitat loss, incidental take from various industry projects, and lethal White-nose Syndrome disease. It is critical to accurately census habitat for appropriate conservation measures, yet traditional sampling methodology, such as mist netting and acoustic recordings, can be time-intensive and biased. Instead, a passive sampling tool that does not rely on the *a priori* knowledge of bat roosts may provide crucial information on bat communities. In the water-limited habitats of forested uplands of the Appalachian Plateau, water-filled road-ruts are important resources for bats. Therefore, we developed an environmental DNA (eDNA) protocol to sample isolated road-ruts that may have the presence of sloughed cellular material from actively drinking bats. The detection of bat eDNA was investigated from a positive control experiment, and across 47 water samples collected in Kentucky and Ohio. Water samples were analyzed using both species-specific quantitative polymerase chain reaction (qPCR) and community metabarcoding methodologies. Using qPCR analysis, we detected eDNA from big brown bat (*Eptesicus fuscus*) and eastern red bat (*Lasiurus borealis*) from water-filled road-ruts. While the community metabarcoding approach failed to detect any bat eDNA, many non-target amphibians, birds, and mammals were identified. These results suggest eDNA found within road-ruts provides an additional detection tool for surveying biodiversity across upland forests. Additionally, the use of qPCR increased the detection of rare eDNA targets, which will be crucial for properly implementing future eDNA applications for improving bat conservation efforts across the landscape.

**Article impact statement:** Environmental DNA provides detection of bats from drinking sources offering a novel survey method for management and conservation efforts

## 1. INTRODUCTION

Bats are the most widely distributed terrestrial mammal on Earth with >1,400 global species, which provide crucial ecosystem services in both natural and agricultural ecosystems (Kunz et al., 2011). However, local biodiversity of North American temperate forest bat populations is rapidly declining due to anthropogenic impacts and habitat loss (Carter 2006, Frick et al. 2017, 2020), as well as the lethal White-nose Syndrome (WNS) disease caused by the invasive fungal pathogen *Pseudogymnoascus destructans* (Frick et al., 2010; O’Keefe et al., 2019). Of the 14 species of bat occurring within the Appalachian Plateau of eastern United States, four are protected and managed under the Endangered Species Act (ESA), including the Virginia big-eared bat (*Corynorhinus townsendii virginianus*), the gray bat (*Myotis grisescens*), the northern long-eared bat (*M. septentrionalis*), and the Indiana bat (*M. sodalis*). In addition, some individual states regulate their own lists of endangered, threatened, and species of special conservation concern (see ODNR, 2020). Additionally, compositional changes in bat communities are occurring throughout North America following the 2006 invasion of WNS, affecting small-bodied bat populations disproportionally to large-bodied bats (Silvis et al., 2016), and resulting in overall declines in *Myotis* spp. and tri-colored bats (*Perimyotis subflavus*) (Pettit & O’Keefe, 2017; Nocera et al., 2019a). Thus, monitoring of both year-round and migratory bat presence and abundances is vital for appropriate population management plans for regulators, such as the mobile acoustic bat monitoring by the United States Fish and Wildlife Service (Evans et al., 2021) and continued hibernacula monitoring every two years.

Species composition and population data traditionally are collected using a combination of methods, such as mist netting (Kiser & MacGregor, 2004; Robbins et al., 2008), colony counts (Warren & Witter, 2002), and acoustic monitoring (Simonis et al., 2020). While mist netting involves simple, inexpensive equipment, it can be time consuming, require expert knowledge on bat behavior and taxonomy (MacCarthy et al., 2006), and poses a risk for WNS transmission across populations (see USFWS, 2016). Colony counts assess occupancy, abundance, and distribution for roosting bats, but require *a priori* knowledge and expertise for finding roosting locations, pose hazards to surveyors (Warren & Witter, 2002), and additionally poses a risk for WNS transmission. Acoustic monitoring overcomes reliance on handling bats directly by detecting ultrasonic calls used by individual bat species, however calls may not definitively distinguish all species and some species display large behavioral plasticity in vocalizations (Barclay, 1999; Jones & Siemers, 2011). Additionally, acoustic surveys favor species that produce low frequency and high intensity calls (Griffin, 1958), and acoustic monitoring software can lead to biased species detections (Samuel et al., 1992; Nocera et al., 2019b). Therefore, the development of new survey methods may circumvent limitations of existing methods for monitoring bat populations

DNA analysis of guano from roosts provides presence/absence and other valuable data, such as diet analysis and the detection of pathogens (Walker et al., 2016, 2019; Swift et al., 2018). Similar to acoustic monitoring, DNA analysis of guano provides a non-invasive survey tool that does not rely on the handling of bats, preventing the concern of spreading disease or the potential risks of injuries to bats and survey staff. However, guano collection requires knowledge of and access to the exact location where the target organism is or was recently present, and is not a passive sampling tool for solitary roosting species. Acquiring a baseline understanding of bat biology and spatial distribution is necessary for the management of this taxonomic group, however documenting many species on a spatial level can be difficult due to behavioral variability (Barclay, 1999; Peixoto et al., 2018), and uncertainty in roosting locations. Instead, a passive sampling tool that does not rely on the *a priori* knowledge of bat roosts may provide crucial information on status and distribution of bat communities.

Major advancements over the past decade through the analysis of environmental DNA (eDNA – genetic material released from urine, waste, mucus, or sloughed cells) have considerably improved surveys for a wide-range of aquatic taxa (Beng & Corlett, 2020; Deiner et al., 2021). More recently, methods for eDNA collection from aquatic environments have been developed for detection of terrestrial mammals, such as coyotes (*Canis latrans*) (Rodgers & Mock, 2015), invasive wild boar (*Sus scrofa*) (Davis et al., 2018), elusive jaguar (*Panthera onca*) (Wilcox et al., 2021), and even entire terrestrial mammal communities (Ushio et al., 2017; Harper et al., 2019). The detection of terrestrial mammals from an aquatic water source is dependent upon the frequency and duration the species spends drinking and/or wading through the body of water (Seeber et al., 2019), and the time since the species last visited (Farrell et al., 2020). Even with the limited release of terrestrial eDNA compared to aquatic fauna, some terrestrial birds (Day et al., 2019) and mammals (Sales et al., 2019) have displayed greater detection probability with eDNA compared to traditional monitoring methods.

As presence/probable absence surveys for bats, especially those protected under ESA and within state regulations, can be time intensive and costly, eDNA sampling from drinking sources may provide an additional survey technique to improve conservation efforts. For bats in particular, traditional survey methods often target riparian zones, river systems, and still bodies of water where individuals gather from surrounding populations (Kiser & MacGregor, 2004; Brooks & Ford, 2005; Johnson et al., 2010), therefore these locations may also be suitable for bat surveys utilizing eDNA. However, the only investigation into aquatic eDNA bat detection found limited detection of big brown bat (*Eptesicus fuscus*) eDNA from streams in western US despite visual confirmation of presence (Serrao et al., 2021). Water samples collected across four rivers yielded only a single positive detection at very low concentrations (i.e., below limit of detection), suggesting eDNA sampling from lotic systems might limit detection of bats as their saliva may quickly become diluted and transported down river. Alternatively, considering small ponds and water-filled road-ruts in forested uplands are important sources of water for bats, (Wilhide et al., 1998; Kiser & MacGregor, 2004), eDNA sampling from these smaller lentic drinking sources may be better suited for bat detection.

An eDNA approach that samples source drinking water offers the potential to survey bat occupancy without reliance on finding guano or the *a priori* knowledge of roost locations. The Electric Power Research Institute (EPRI) and American Electric Power (AEP) had previously commissioned research describing bat detections using mist netting and acoustic recordings, and expressed new interest in the efficacy of eDNA methods. Therefore, the objective of this study was to test for the detection of bat eDNA from road-ruts in mixed-mesophytic forests spanning across Ohio and Kentucky during spring and summer months. This study provides an important resource for future conservation efforts and monitoring of threatened bat populations by providing a supplementary tool for documenting presence of a species.

## 2. METHODS

### 2.1 Control Bat Dip (CBD) Positive Control

To gain an understanding of possible bat eDNA release from a drinking occurrence within a road-rut, we developed an experiment to simulate a bat drinking event within a large Rubbermaid tub of distilled water. This ‘Control Bat Dip’ (CBD) was conducted on August 19, 2020 during the collection of the environmental samples from site KY-S3 (Figure 1, see section 2.2.3). Bats were collected from a nearby mist net and handled by experienced and trained bat ecologists following the Kentucky Department of Fish and Wildlife Resources Scientific Wildlife Collecting Permit (SC2011194). Bats were handled while wearing new latex gloves between each individual. Two net sets were erected adjacent to a single water-filled road-rut (not associated with KY-S3). This CBD sample consisted of “simulating” drinking of bats for two big brown bats (14.7g and 19g), one eastern red bat (14.5g), and one eastern small-footed bat (*Myotis leibii*) (5.2g). Each individual bat was dipped six times in a motion that mimics drinking, with their mouth/face, chest, and stomach grazing the surface of the water. The CBD was performed in 4.5 liters of distilled water in a 34 liter Rubbermaid tub. Following the complete dipping of each of the four individual bats, two 1 liter water samples were collected in sterile 1 liter Nalgene bottles. The water was filtered via vacuum-pump from one sample using a 47-mm-diameter polycarbonate filter (PC, pore size 2 µm; Cytivia Whatman, United Kingdom) and the other using a 55-mm-diameter glass microfiber filter (GF, nominal pore size 11 µm; Cytivia Whatman, United Kingdom). After vacuum-pumping, filters were placed in 15ml vials filled with RNALater and stored/shipped on ice to the Northern Arizona University (NAU) Bat Ecology & Genetics Laboratory for analysis.

**Figure 1.**
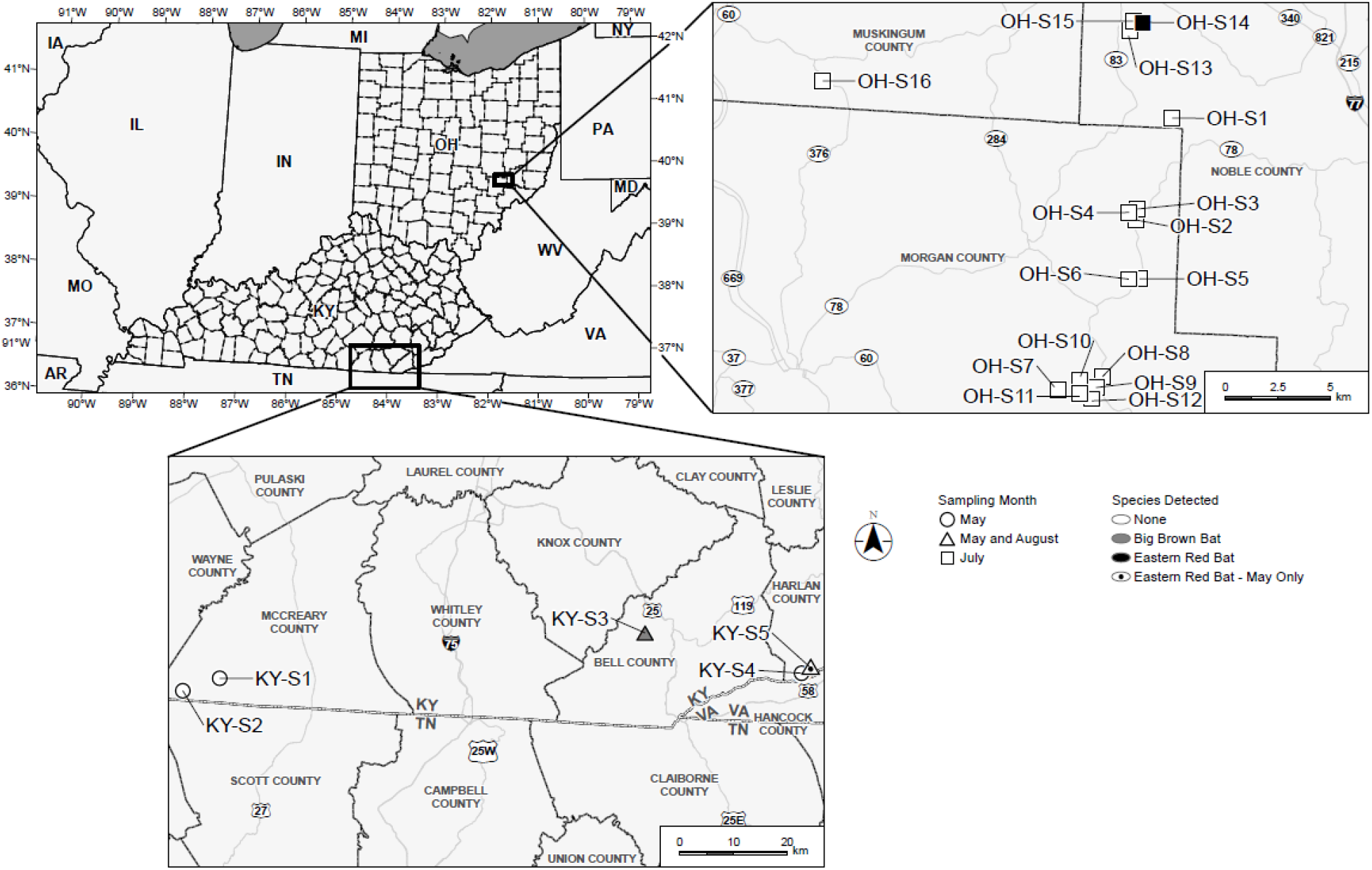
Sampling locations of water-filled road-ruts in Kentucky and Ohio. Month of sampling is labeled by shape (circle = May, square = July, and triangle = May and August), and filled shapes are labeled as bat detection (grey = big brown bat (*Eptesicus fuscus*) and black = eastern red bat (*Lasiurus borealis*)).

### 2.2 Survey sites and eDNA collection

Environmental DNA water samples were collected from 47 samples across 21 road-ruts from three counties in Kentucky and three counties in Ohio sampled across May, July, and August of 2020 (Figure 1, Supplementary File 1). Each set of samples are detailed below.

#### 2.2.1 May 2020 – Kentucky Field Sampling

The initial field sampling occurred between May 23 – 29, 2020 at five sites in Kentucky across Bell, Harlan, and McCreary Counties (Figure 1). Three sites contained tire track linear road-ruts with standing water in the deepest portions, while two sites contained circular puddles of water in low spots. These water sources were sampled using a sterile 1 liter Nalgene bottle, skimming the water off the surface with a gloved hand. Water was filtered through single use analytical filter funnels with a 47mm diameter cellulose nitrate membrane (CN, pore size 0.2 µm; Thermo Fisher Scientific) using a drill powered inline pump. One liter was filtered through each filter (Supplementary File 1). Filters were placed in 15ml vials filled with RNALater and stored/shipped on ice to NAU for analysis.

#### 2.2.2. July 2020 – Ohio Field Sampling

The second round of field collection occurred on July 28 – 29, 2020 on the former AEP ReCreation Land property which spans Noble, Morgan, and Muskingum Counties in Ohio (Figure 1). Sites were chosen based on an experienced biologist’s visual assessment of their ability to retain water, or remain perennial throughout the maternity and post-maternity season (June – August) when lactating female bats and juveniles require more water. These sites varied from large circular pools (>100 square meter), small circular pools (<100 square meter), potholes, and linear truck tire ruts. At each site, two samples were taken in sterile 1 liter Nalgene bottles by skimming the top of the water body. The water was filtered via vacuum-pump from one sample using a 47-mm-diameter polycarbonate filter (PC pore size 2 µm; Cytivia Whatman, United Kingdom) and the other using a 55-mm-diameter glass microfiber filter (GF nominal pore size 11µm; Cytivia Whatman, United Kingdom). Between one and three polycarbonate filters were combined into an aggregate sample to attempt to filter at least 50ml of water per road-rut. At two sites (OH-S3 and OH-S11), two additional 1 liter samples were taken after disturbing the substrate with a gloved hand, to suspend any settled material back into the water column prior to sampling. Filtration of samples ranged from 25–950ml for the 16 sampling sites (Supplementary File 1). Filters were preserved as listed above.

#### 2.2.3 August 2020 – Kentucky Field Sampling

The third round of sampling occurred on August 19 – 20, 2020 in Kentucky at two of the sites previously sampled in May, (KY-S3 and KY-S5, Figure 1). Water samples were collected from the two sites at the beginning of the night, after viewing bats actively drinking, and at the end of the net night with the filtration methods described above (section 2.2.2). Additionally, samples taken at the end of the net night were first taken without disturbing the water pool or underlying sediment, with an additional sample taken after disturbing the water column and sediment interface to resuspend any DNA material that may have settled. Water was processed with the filtration methods described above (section 2.2.2), with the amount of filtrate varying from 25–125ml through the PC filters and 175–500ml through the GF filters (Supplementary File 1). Filters were preserved as listed above.

### 2.3 eDNA analysis

We tested for bat eDNA detection using both community metabarcoding and species-specific quantitative polymerase chain reaction (qPCR) methodologies. These two methodologies differ in their laboratory processes, and in their detection limits (Bylemans et al., 2019).

#### 2.3.1 High-throughput sequencing metabarcoding

The DNA from each filter was extracted by NAU using a modified Qiagen DNeasy® Blood and Tissue Kit (Qiagen Inc.) protocol (Walker et al., 2016, 2019). Environmental samples collected from Kentucky sites in May 2020 were analyzed with metabarcoding targeting a ∼202 base pair (bp) fragment of the mitochondrial (mtDNA) cytochrome oxidase subunit I (COI) gene that has been specific for bats in guano samples (Walker et al., 2016, 2019). Samples collected in July and August 2020 from Ohio and Kentucky, respectively, were additionally analyzed with metabarcoding targeting a ∼100 bp fragment of the mtDNA 12S gene that is universal to vertebrates (Riaz et al., 2011). Library preparation consisted of a 2-step PCR, in which the first step utilized the target marker (COI or 12S), followed by a second reaction with incorporated universal tails (UT1 on 5’ end of forward primer: ACCCAACTGAATGGAGC; UT2 to 5’ end of reverse primer: ACGCACTTGACTTGTCTTC). The first PCR contained 2 μL undiluted DNA template in a 10 μL reaction, with 1 μL 10X Mg-free PCR buffer (Invitrogen, Thermo Fisher Scientific, Waltham, MA, USA), 2.5 mM MgCl2, 0.2 mM of each dNTP, 0.4 μM primers, and 0.3 U/ μL PlatinumTaq DNA polymerase (Invitrogen, Thermo Fisher Scientific, Waltham, MA, USA). Cycling involved an initial step of 95°C for 10 min, followed by 38 cycles of 60 s at 95°C, 30 s at 60°C, and 30 s at 72°C, and concluding with a final extension step of 72°C for 10 min. The resulting amplicons from this initial PCR were employed as template in a subsequent Illumina (MiSeq) extension PCR using unique Illumina indices containing sequences complementary to the universal tails. The second PCR incorporating the UT-labeled primers was scaled up to a 20 μL reaction with 1 μL of undiluted PCR product from the first step, 0.1 U/μL Platinum Taq, and 35 cycles. A no template control (NTC) was processed with each set of PCRs.

Library preparation used a dual indexing strategy, with each sample tagged by two 12 nucleotide-long indices each at 3–4 mismatches away from every other index in the set. This results in a combined index length of 24 nucleotides and 6–8 mismatches from any other dual index, which greatly limits the amount of index hopping (Walker et al., 2019). For each experiment, libraries were quantified using KAPA Illumina Library Quantification Kit (KK4933) and indexed samples were pooled in equal concentration. We sequenced pools for each experiment using Illumina MiSeq V3 600 cycle kit as flow-cell space became available. These runs also included samples that were not part of the current project.

Sequencing reads were computationally processed to obtain the highest quality taxonomic results in QIIME2 v2020.2 (Bolyen et al., 2018). Priming regions, adapters, and read-through were removed using cutadapt v2.1 (Martin, 2011) to isolate the fragments of interest. We removed low quality reads, alleviated sequencing contamination by joining paired-end reads, filtered out PCR artifacts (chimeric reads) using DADA2 (Callahan et al. 2016), and subsequently filtered improperly joined reads. Using a positive control consisting of DNA from known bat species, spotted bat (*Euderma maculatum*) and Townsend’s big-eared bat (*C. t. townsendii*), we identified a read threshold by which to filter out read variants of likely sequencing error, allowing us to retain 99.5% reads/sample. Sequences for COI were then classified using a naïve-Bayes machine learning classifier (Bokulich et al., 2018) that was trained against a custom, curated, reference database of bat species. We retained species classifications only if they were classified by at least 90% bootstrap support. Read variants of the 12S marker and any read variants of COI not classified using the machine learning algorithm were cross-referenced against the National Center for Biotechnology Information’s (NCBI) GenBank database (Benson et al., 2009) using BLAST (Altschul et al., 1990) and taxa classified from those results using Least Common Ancestor (LCA) analysis in MEGAN v6 (Huson et al., 2007).

#### 2.3.2 Quantitative PCR

Extracted DNA samples were sent to Cramer Fish Science Genidaqs (https://genidaqs.com) for qPCR analysis. The primer probe set for big brown bat was previously developed and tested for specificity for a separate bat-related project at Stantec with Genidaqs. Genidaqs further developed a new primer probe set for the eastern red bat from samples provided by Dr. Joseph Johnson’s Laboratory at Ohio University. Primer and probe optimization were conducted following Applied Biosystems (Thermo Fisher ABI) guidelines for optimizing primer and probes for amplifying custom target sequences. PCR for optimization was performed in 10 μl total volume containing 4 μl of DNA template, 5 μl TaqMan™ Environmental Master Mix (Thermo Fisher ABI), 0.5 μl each, 50-900 nM final concentration of both forward and reverse primers, and 1 μl 50–250 nM final concentration probe. Thermocycling of PCR reactions were conducted on a QuantStudio™ 3 Real-Time PCR System (Thermo Fisher ABI) with the following cycle conditions: initial activation 10 min at 95° C followed by 40 cycles of 15 sec denaturation at 95° C and 1 min extension at 60° C. The newly developed eastern red bat assay was tested for specificity *in-vitro* using DNA as a template from vouchered specimens of eastern red bat and closely related and potentially co-existing congeners (big brown bat, silver-haired bat (*Lasionycteris noctivagans*)). The PCR for specificity was performed in triplicate in 10 μl total volume containing 4 μl of 2 ng/μL of genomic DNA template, 5 μl TaqMan Universal Master Mix (Thermo Fisher ABI), 0.5 μl each, 900 nM initial concentration of both forward and reverse primers and 0.2 μl of either 10 μM big brown bat or 5 μM eastern red bat initial probe concentration. All PCR reactions were conducted with three NTC reactions, to assess for sample contamination.

To determine the sensitivity of the assay, tenfold serial dilutions of genomic DNA ranging from 0.0000002 ng/μL to 2 ng/μL was amplified using each assay following the protocol listed above. Each of the tenfold serial dilutions was amplified eight times along with four no template controls using the same cycle conditions as described for specificity and optimization. Quantitative PCR assays were evaluated for sensitivity based on the qPCR efficiency, limit of quantification (LOQ) – defined as the lowest concentration of target that can be accurately quantified with a coefficient of variance below a threshold of ≤35%, and limit of detection (LOD) – defined as the lowest concentration of DNA that can be detected in 95% of replicates (see Klymus et al., 2020).

Each environmental sample was analyzed with both bat assays in sextuplet (with a few exceptions based on limited extraction volume), with each qPCR technical replicate consisting of a 10 μl reaction volume. Each 10 μl qPCR reaction was composed of 5 μl TaqMan Universal Master Mix (Thermo Fisher ABI), 0.1 μl each, 900 nM initial concentration of both forward and reverse primers and either 0.2 μl 10 μM big brown bat or 5 um eastern red bat initial concentration probe and 4 μl DNA template. Thermocycling protocol followed that listed above. Six NTC reactions were run on the plate with the control sample templates consisting of 4 μl of ultrapure water replacing DNA template within reaction volume. Three positive control reactions consisting of 2 ng/μL of target species genomic DNA as template were also tested in parallel to ensure consistent PCR performance.

## 3. RESULTS

### 3.1 Quantitative PCR assay design

One qPCR assay was developed and tested for specificity of eastern red bat, however it displayed a potential for cross-amplification with DNA from silver-haired bat. Silver-haired bat worked poorly as demonstrated by a three-fold reduction in quantification of known starting concentration. For example, silver-haired bat at a concentration of 2 ng/ul quantified as 0.02 ng/ul with the eastern red bat primers and probe. Although this is a substantially lowered detection, silver-haired bat must be considered as potentially cross-amplifying, and therefore, any positive detection with this qPCR assay may indicate the presence of either species within a sample. Considering our sampling sites, historical bat presence in the study area during sample collection period, and recent mist netting data, we determined the likelihood of silver-haired bat presence to be very low, and thus any positive detections for this assay are reported here as eastern red bat. The big brown bat assay was designed for the eastern haplogroup and may present lower efficiency with the western haplogroup. Previously, Serrao et al. (2021) developed big brown bat assays that are capable of discerning haplogroup population variation. The big brown bat assay displayed a qPCR efficiency of 93.0%, with a LOQ of 2*10^−4^ ng of genomic DNA and a LOD of 2*10^−5^ ng of genomic DNA. The eastern red bat assay displayed a qPCR efficiency of 94.3%, with a LOQ of 2*10^−5^ ng of genomic DNA and a LOD of 2*10^−6^ ng of genomic DNA. The high qPCR efficiency suggests both assays perform well and can accurately quantify bat DNA at a range of concentrations.

### 3.2 Control Bat Dip (CBD)

#### 3.2.1 CBD: High-throughput sequencing metabarcoding

The PC filters successfully detected bats for both the COI and 12S assays, while the GF filter only successfully detected bats for the 12S marker (Figure 2). The GF filter with the 12S marker was the only sample to discern all three bat species (including eastern small-footed bat), with the PC filters only detecting big brown bat and eastern red bat (Figure 2). Of bat detections, the majority of reads were of big brown bat on both filter types (PC COI: 97.94%, PC 12S: 96.64%, GF 12S: 67.81%) (Figure 2). The COI assay was able to provide greater taxonomic resolution than the 12S assay, as 12S was only able to discern to the genus-level, while COI went to species level. The control samples also contained large amounts of non-target contaminated sources, such as reads corresponding to human, dog, and cat (PC 12S: 64.84%, PC COI: 54.56%, GF COI: 51.36%).

**Figure 2.**
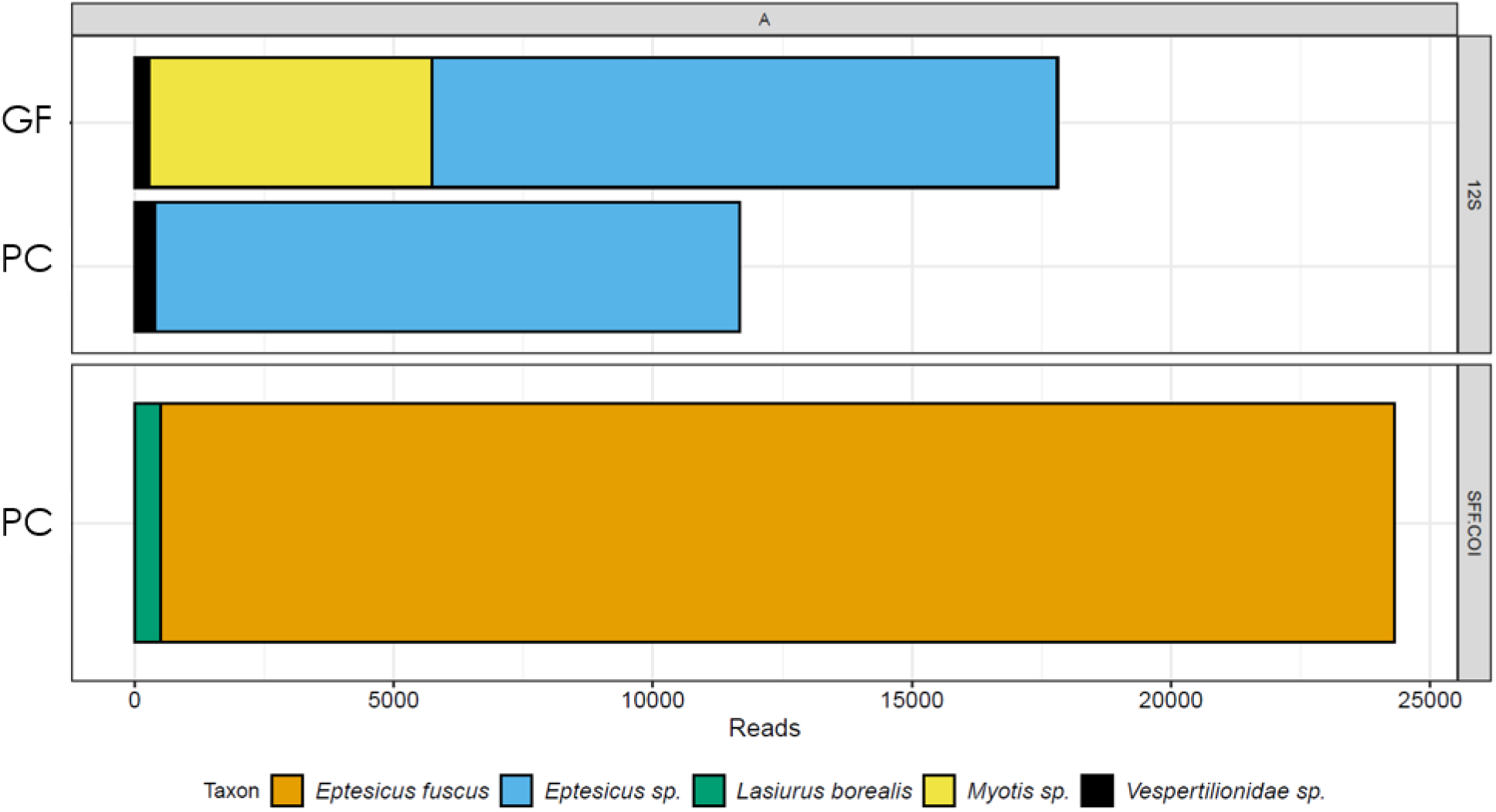
Number of high-throughput sequence reads and their species identity from the control bat dip (CBD) using two filter types (Glass microfiber 11µm pore (GF) and Polycarbonate 2µm pore (PC)) and targeting a ∼100 base pair (bp) fragment of the mitochondrial (mtDNA) 12S gene that is universal to vertebrates (Riaz et al., 2011) and ∼202 bp fragment of the mtDNA cytochrome oxidase subunit I (COI) gene developed for bats (Walker et al., 2016, 2019). Note the *Myotis sp*. represents the eastern small-footed bat (*Myotis leibii*).

#### 3.2.2 CBD: Quantitative PCR

Quantitative PCR successfully detected big brown bat and eastern red bat eDNA in the control bat dip sample for both the PC and GF filters. While both filter types successfully detected the target bat species, there were clear differences in the eDNA concentration and detection rate based on filter type. GF displayed greater DNA concentration for both big brown bat (GF 311.21±57.85 (fg/reaction); PC 21.78±12.37) and eastern red bat (GF 5.47±1.02 (fg/reaction); PC 2.21) (Table 1 and Figure 3). Additionally, for eastern red bat, only one out of three technical replicates was positively detected for PC, while all three technical replicates were positive for GF. Furthermore, for both species, GF displayed a higher number of qPCR replicates above the LOQ and LOD (Table 1 and Figure 3). When comparing concentrations of eDNA between bat species, big brown bat displayed greater DNA concentration compared to eastern red bat for both filter types (Figure 3).

**Table 1.** Quantitative PCR (qPCR) results from control bat dip samples for big brown bat (*Eptesicus fuscus*) and eastern red bat (*Lasiurus borealis*). Positive detections are listed as the mean quantification cycle (Cq value) ± standard deviation and the quantified concentration of genomic (g)DNA ± standard deviation. The number of positive replicates, the number of replicates above the limit of quantification (>LOQ) and the limit of detection (>LOD) are additionally listed.

| Species | Filter Type (Pore Size) | Cq ± SD | Concentration ± SD (fg/μL) | N | >LOQ (N) | >LOD (N) |
| --- | --- | --- | --- | --- | --- | --- |
| Big brown bat | PC (2.0 μm) | 35.71±0.79 | 21.78±12.37 | 3 | 0 | 1 |
| | GF (11 μm) | 31.54±0.28 | 311.21±57.85 | 3 | 3 | 3 |
| Eastern red bat | PC (2.0 μm) | 37.16 | 2.21 | 1 | 0 | 1 |
| | GF (11 μm) | 35.81±0.27 | 5.47±1.02 | 3 | 0 | 3 |

**Figure 3.**
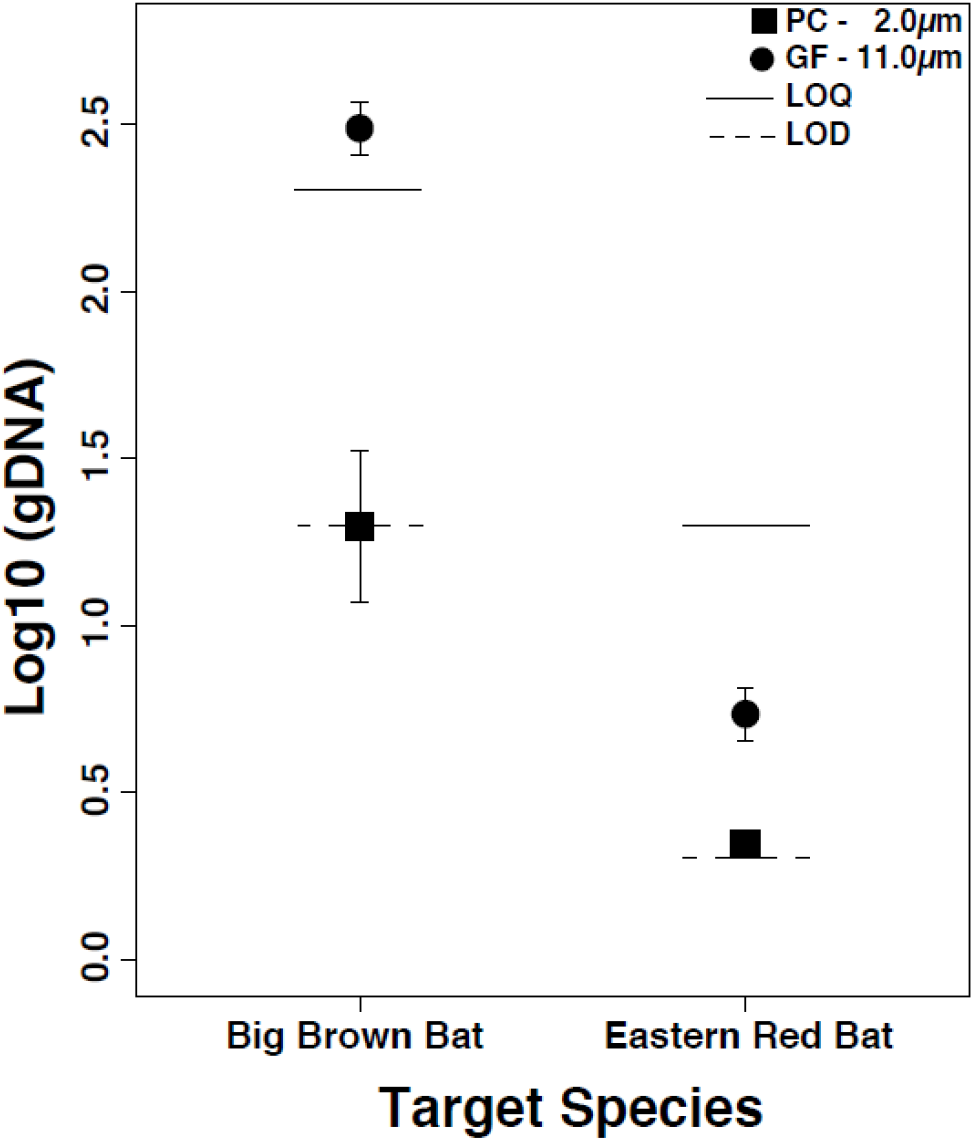
The mean genomic (g)DNA concertation (log10 of femtograms per µL) from quantitative PCR (qPCR) for big brown bat (*Eptesicus fuscus*) and eastern red bat (*Lasiurus borealis*) from control bat dip (CBD) samples that were filtered on polycarbonate (PC 2µm) or glass microfiber (GF 11µm) filters. Error bars represent standard deviation across three qPCR replicates. Note only one of three qPCR technical replicates detected DNA for the PC 2µm filter for eastern red bat. The limit of quantification (LOQ) and limit of detection (LOD) are depicted for each species-targeted assay.

### 3.3 Environmental samples

#### 3.3.1 Environmental samples: High-throughput sequencing metabarcoding

There were no signs of laboratory contamination, as all NTC samples contained no vertebrate sequence reads. Of the environmental samples, 36 successfully sequenced for the 12S marker and 27 for the COI marker. For the 12S marker, after sequence processing samples ranged from 4,505–77,403 sequence reads (mean 33,774.16 ± 16,410.14), retaining on average 87.19 ± 12.03% of sequence reads. For the COI marker, after sequence processing samples ranged from 10,017–53,607 sequence reads (mean 27,719.69 ± 10,417.13), retaining on average 48.10 ± 20.96% of sequence reads. No bat sequences were detected from metabarcoding in any of the 47 environmental samples. After removing human related read sequences (e.g., human, dog, cat, pig, chicken, and cow), 16 total non-bat vertebrate taxa were detected within the environmental samples across both markers (10 species detected with 12S and nine species detected with COI). Based on known species distributions we conclude the presence of the following species with the 12S marker: amphibians (*Anaxyrus sp*. (American toad or Fowler’s toad), *Hylidae sp*. (tree frogs), *Rana clamitans* (green frog), and *Rana sylvatica* (wood frog)), birds (*Passeriformes sp*. (possible Turdidae)), and mammals (*Didelphidae sp*. (Virginia opossum), *Odocoileus sp*. (white-tailed deer), *Prycon sp*. (racoon), and *Peromyscus sp*. (deer mice)) (Table S1). Species detected with the COI marker include amphibians (*Anaxyrus sp*., *Hylidae sp*., green frog, wood frog, and *Notophthalmus viridescens* (eastern newt)), birds (*Sayornis phoebe* (eastern phoebe) and *Hylocichla mustelina* (wood thrush)), and mammals (*Lynx rufus* (bobcat) and *Ursus americanus* (black bear)) (Table S1).

#### 3.3.2 Environmental samples: Quantitative PCR

Of the 47 samples analyzed, qPCR detected big brown bat in two samples collected from Kentucky from the same sampling location (KY-S3) collected in May and August (Figure 1, Table 2). Both positive samples for big brown bat were below the LOQ, with only the sample collected in August displaying concentrations above the LOD (Table 2). However, even with low concentrations, both samples displayed positive detections in multiple technical replicates (50% positive in May (3/6) and 67% positive in August (2/3)) (Table 2). Eastern red bat was positively detected in one sample from Kentucky (KY-S4) on 05/29/2020 and one sample from Ohio (OH-S14) on 07/29/2020 (Figure 1, Table 2). Detections for eastern red bat displayed very minute concentrations of bat eDNA, with all positive samples displaying values below LOQ and LOD (Table 2). While the KY-S4 sample displayed positive in 83% of technical replicates (5/6), the OH-S14 was only positive in 17% of technical replicates (1/6) (Table 2). Across the four positive environmental samples, 75% (3/4) corresponded to samples with 1000 mL of filtrate (Table 2). Positive detections occurred across all three sampling months and with all three filter matrices/pore sizes (Table 2).

**Table 2.**
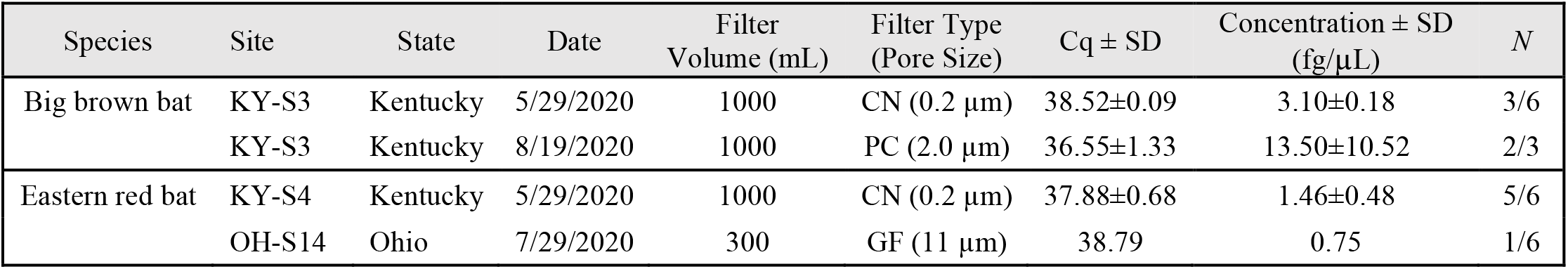
Environmental DNA samples with positive detections from quantitative PCR (qPCR) for big brown bat (*Eptesicus fuscus*) and eastern red bat (*Lasiurus borealis*) collected from Kentucky and Ohio upland forests. Positive detections are listed as the mean quantification cycle (Cq value) ± standard deviation and the quantified concentration of genomic (g)DNA ± standard deviation. The number of positive qPCR replicates is additionally listed.

| Species | Site | State | Date | Filter<br>Volume (mL) | Filter Type<br>(Pore Size) | Cq ± SD | Concentration ± SD<br>(fg/μL) | N |
| --- | --- | --- | --- | --- | --- | --- | --- | --- |
| Big brown bat | KY-S3 | Kentucky | 5/29/2020 | 1000 | CN (0.2 μm) | 38.52±0.09 | 3.10±0.18 | 3/6 |
| | KY-S3 | Kentucky | 8/19/2020 | 1000 | PC (2.0 μm) | 36.55±1.33 | 13.50±10.52 | 2/3 |
| Eastern red bat | KY-S4 | Kentucky | 5/29/2020 | 1000 | CN (0.2 μm) | 37.88±0.68 | 1.46±0.48 | 5/6 |
| | OH-S14 | Ohio | 7/29/2020 | 300 | GF (11 μm) | 38.79 | 0.75 | 1/6 |

## 4. DISCUSSION

Environmental DNA sampling continues to expand beyond identification of aquatic taxa (Harper et al., 2019), and new methods have begun to show promise for assessing the presence of terrestrial organisms. Here, we design a unique sampling strategy targeting small water-filled road-ruts in upland forests, to collect eDNA from drinking water sources utilized by bats (Kiser & MacGregor, 2004). In many areas of upland forests in the Appalachian Plateau of eastern United States, these road-ruts created by 4-wheel drive and all-terrain vehicles are the closest source of drinking water for some bat populations, especially near ridgetops where solar-exposed maternity roost trees are most likely to occur (Kiser & MacGregor, 2004). We successfully detected bat eDNA from four road-ruts within mixed-mesophytic forests at three widely separated locations within Ohio and Kentucky. This is only the second documentation of bat detection from aquatic eDNA, including the first documentation for eastern red bat. These results suggest the small volumes of flooded road-ruts potentially contain detectable concentrations of bat eDNA, providing a complementary methodology to use alongside acoustic and mist netting techniques for determining presence/probable absence of bats within upland forests.

Within the experimental CBD samples, both species-specific qPCR assays successfully detected their target species (big brown bat and eastern red bat) across both filter types, and metabarcoding additionally detected eastern small-footed bat with the GF filter. Big brown bat displayed greater eDNA concentrations compared to eastern red bat, suggesting the higher number of individuals used and/or the larger size of big brown bat equated to higher eDNA shedding rates in the CBD samples. However, in a natural environment, inferences on bat abundance from eDNA quantification may be limited (see review in Yates et al., 2019). Several bat species display variations in their molting cycle (Fraser et al., 2013) and time of nursing their young, which are likely to cause seasonal fluctuations in rates of fur shedding and drinking frequencies (Daniel et al., 2010). Bat eDNA surveys may be better suited for presence/probable absence-based surveys, to provide a quick and inexpensive screening tool for depicting sites that can benefit from future mist net surveys. The CBD experiment detected higher concentrations of eDNA on the GF filters compared to PC filters for both target species, suggesting the GF matrix was more successful in retaining bat eDNA. GF filters are the most frequently used filter for aquatic eDNA capture and have been recommended for fish eDNA collection (see review in Shu et al., 2020). The larger pore size of the GF filter (11μm vs 2μm) did not appear to negatively impact eDNA retention, which likely corresponds to the eDNA in the CBD samples originating from intact cells and tissues of saliva and hair follicles.

The mean bat eDNA concentration was greater within all of the CBD samples compared to any of the environmental samples. While the setup attempted to mimic the act of bats drinking by skimming their bellies across the top of the water and potentially releasing hair and salvia, the CBD may have released greater amounts of epithelial cells and hair follicles compared to the natural act of bats drinking. Furthermore, the CBD was also performed in a small volume of water (4.5 liters) compared to the environmental samples, and consisted of distilled water opposed to the high sedimentation and organic content present in some environmental samples. These factors likely influenced the higher concentrations found within the CBD samples. Although their control setup differed from the current study, Serrao et al. (2021) likewise found greater concentrations of bat eDNA in control samples compared to concentrations in the natural environment.

Using qPCR, we detected big brown bat in two environmental samples from Kentucky and eastern red bat from one sample in Ohio and one sample in Kentucky. Similar to Serrao et al. (2021), we found low detection rates and low eDNA concentrations for big brown bat within an aquatic environment. In our case, all positive samples fell below LOQ and all but one below LOD. Alternatively, with a metabarcoding approach we failed to detect any bat species with the combination of a 12S and a COI metabarcoding marker. Furthermore, within the CBD samples we were only able to successfully detect all three bat species with the 12S metabarcoding marker on the GF filter, while the PC filter failed to detect eastern small-footed bat with either metabarcoding marker. These results suggest the metabarcoding markers used were less sensitive for the detection of the low concentrations of bat eDNA present within the samples. Similarly, previous studies have determined higher sensitivity for eDNA detection of aquatic organisms with qPCR compared to metabarcoding methods (Bylemans et al., 2019).

While we failed to detect bat species in the road-ruts using metabarcoding, we did detect a large amount of biodiversity from non-target mammals, birds, and amphibians. The COI metabarcoding assay used in the current study have previously been successful in metabarcoding bat guano samples (Walker et al., 2016, 2019); however, the low bat eDNA concentrations found within the road-ruts appear to prevent appropriate detection when found mixed with a high abundance of other vertebrate non-target eDNA (e.g., amphibian). Metabarcoding allows the additional identification of non-target taxa that co-amplify with the utilized assay, although these non-target DNA fragments can greatly reduce the ability to detect target DNA when it is rare (Marshall & Stepien, 2020). The breeding season of non-target amphibians may reduce bat eDNA detection as the amphibian larvae may directly consume it, and the increased concentrations of amphibian eDNA due to spawning and larval activity may reduce its amplification with the current metabarcoding markers. Designing bat-specific assays or utilizing a mammal-specific assay (Ushio et al., 2017; Sales et al., 2019) might provide better sensitivity for bat eDNA metabarcoding.

The majority of the environmental samples with positive bat detections correspond to filters with the greatest volume (1000 mL), suggesting larger volumes may be required for improved detection rates. Furthermore, biological replicates have been shown to be important in increasing detection probability of rare eDNA targets (Tingley et al., 2021), and thus future bat eDNA surveys should include replicate sampling in the study design. Additionally, multiple environmental factors and site characteristics will likely influence the ability to successfully detect bat eDNA. For example, smaller bodies of water may provide higher detection rates due to lower eDNA dilution (Seeber et al., 2019), while open canopy water bodies may reduce eDNA detection (Day et al., 2019) due to an increase in ultraviolet light related eDNA degradation (Stickler et al., 2015). The sampling season may also impact concentrations of bat eDNA as the lactation portion of the maternity season (June to August) increases the usage of free-standing water by females and recently volant juveniles (Daniel et al., 2010). However, environmental covariates impacting drinking water selection likely differ between bat species (Francl, 2008). Bats also display differing drinking behaviors, ranging from species dipping entire bodies through the water to some species angling only their face towards the water (J. Kiser pers. observation). These behaviors likely result in differences in the concentrations of hair and tissue material shed into a road-rut. Future analysis utilizing eDNA occupancy modeling may provide a comprehensive understanding of these factors and their influence on bat eDNA detection (Davis et al., 2018), thereby offering a baseline for managers to design opportunistic sampling strategies.

In the mixed mesophytic forests of the Appalachian Plateau of the eastern United States, many state and federal agencies design wildlife water holes in strategic locations to maximize wildlife benefits for a wide range of taxa, including bats (Taylor et al., 2020; KDFWR, 2021). Therefore, these constructed water holes and/or the accidental pools provide rare opportunities to measure terrestrial biodiversity. Novel applications of eDNA techniques may expand our understanding of presence/probable absence and provide new avenues for understanding the reproductive biology and behavior of many elusive animals. Seasonal eDNA monitoring can add a beneficial survey tool for understanding the temporal and migratory patterns of bat populations, and the community-shifts associated with anthropogenic and WNS impacts. Additionally, as wetlands are becoming more recognized as ecologically important bat habitat (Mas et al., 2021), eDNA may be a beneficial monitoring tool to understand the extent of bat occurrence within wetland habitat. While bat eDNA has been collected from sediment (Walker et al., 2019; Serrao et al., 2021), guano (Walker et al., 2016, 2019), and air (Serrao et al., 2021), targeting eDNA from these isolated water holes provides opportunities for detection without *a priori* knowledge of bat roosts. Furthermore, road-ruts may provide the only DNA source for some species that roost singularly in trees, rather than in large colonies. Prior to the arrival of the lethal WNS disease, the now threatened northern long-eared bat represented the most frequently captured species of bat using water sources within the eastern United States (Wilhide et al., 1998; Huie, 2002), and thus road-rut eDNA may add complementary conservation information for surveying and managing these declining populations.

## Acknowledgements

We would like to thank Becca Madsen, Jon Black, and Christian Newman from EPRI for suggesting the addition of an eDNA sampling component to an ongoing study and for their support with the development and implementation of eDNA species monitoring. We would also like to thank AEP for their support and providing access to their lands, specifically Timothy Lohner, John Van Hassel, Jeffrey Wilson, and Tracy Simons. If not for the assistance of Dr. Joseph Johnson at the Ohio University Bat Lab, we wouldn’t have been able to develop the eastern red bat assay, so we would like to thank them for the specimens used for this effort. We would like to thank Dan Godec, Cody Fleece, Josh Adams, Mary Murdoch, Jake Riley, Nathan Noland, and Aaron Kwolek from Stantec for helping with organizing the project, answering questions related to eDNA, reviewing draft manuscript, and collecting samples in the field. Thanks to Jordyn R. Upton for assisting with eDNA metabarcoding library preparation. Finally, we would like to thank Cyndol Kiser, Celya Kiser, Michaela Rogers (Kentucky Department of Fish and Wildlife Resources), Courtney Hayes (Office of Kentucky Nature Preserves), and Robert Myers (Kentucky State Parks) for assisting with the collection and processing of water samples.

---

**Table S1.** Environmental DNA samples from Kentucky and Ohio upland forests with positive detections for non-target vertebrate taxa eDNA with high-throughput metabarcoding sequencing using a ∼100 base pair (bp) fragment of the mitochondrial (mtDNA) 12S gene that is universal to vertebrates (Riaz et al., 2011) and a ∼202 bp fragment of the mtDNA cytochrome oxidase subunit I (COI) gene for bats (Walker et al., 2016, 2019).

| Group | Species | Common Name | 12S | COI |
| --- | --- | --- | --- | --- |
| Amphibians | *Anaxyrus sp.* | American toad, Fowler's toad | OH-S3, KY-S3 | OH-S3, KY-S5 |
|  | *Hylidae sp.* | tree frogs | OH-S6, OH-S7, OH-S9 | OH-S6, OH-S9 |
|  | *Rana sylvatica* | wood frog | KY-S5 | KY-S2, KY-S5 |
|  | *Rana clamitans* | green frog | OH-S3, OH-S11, OH-S13, KY-S5 | OH-S9, OH-S11, OH-S15 |
|  | *Notophthalmus viridescens* | eastern newt |  | KY-S2 |
| Birds | *Sayornis phoebe* | eastern phoebe |  | KY-S2 |
|  | *Hylocichla mustelina* | wood thrush |  | OH-S15 |
|  | *Passeriformes sp.* | possible Turdidae | OH-S15 |  |
| Mammals | *Didelphidae sp.* | Virginia opossum | OH-S14 |  |
|  | *Odocoileus sp.* | white-tailed deer | OH-S3 |  |
|  | *Prycon sp.* | raccoon | OH-S14 |  |
|  | *Peromyscus sp.* | deer mice | KY-S5 |  |
|  | *Lynx rufus* | bobcat |  | KY-S5 |
|  | *Ursus americanus* | black bear |  | KY-S5 |

